# *Sir2* is essential for intergenerational effects of parental diet on offspring metabolism in *Drosophila*

**DOI:** 10.1101/641076

**Authors:** Ryoya Hayashi, Satomi Takeo, Toshiro Aigaki

## Abstract

Recent studies have revealed that parental diet can affect offspring metabolism and longevity in *Drosophila*. However, the underlying mechanisms are still unknown. Here we demonstrate that *Sir2* encoding an NAD+-dependent histone deacetylase is required for the intergenerational effects of low nutrition diet (1:5 dilution of standard diet). We observed an increased amount of triacylglyceride (TAG) in the offspring when fathers were maintained on a low nutrition diet for 2 days. The offspring had increased levels of metabolites of glycolysis and TCA cycle, the primary energy producing pathways. We found that *Sir2* mutant fathers showed no intergenerational effects. RNAi-mediated knockdown of *Sir2* in the fat body was sufficient to mimic the *Sir2* mutant phenotype, and the phenotype was rescued by transgenic expression of wild-type *Sir2* in the fat body. Interestingly, even fathers had no experience of low nutrition diet, overexpression of *Sir2* in their fat bodies induced a high level of TAG in the offspring. These findings indicated that *Sir2* is essential in the fat body of fathers to induce intergenerational effects of low nutrition diet.

## Introduction

Nutritional conditions of parents have significant impacts on offspring growth and development. Recent studies have reported maternal and paternal inter/transgenerational phenotypes, including stress-induced epigenetic changes (Seong *et al.*, 2011), endocrine disruptors (Anway *et al.*, 2016), behavior (Dias & Ressler, 2014), and energy metabolism (Ng *et al*., 2010). Inter/transgenerational inheritance of environment-induced physiology involves generation and transmission of that information to the next generation (Sharma, 2015). It has been shown that altered DNA methylation (Radford *et al*., 2014), histone modifications (Inoue *et al*., 2017; Zenk *et al*., 2017), and noncoding RNA transcripts (Chen *et al.*, 2016) can be transmitted from parents to offspring. However, it is largely unknown how environmental stimuli generate signals leading to inheritable epigenetic modifications.

Transgenerational effects of nutritional conditions have been studied in flies and explored the underlying epigenetic mechanisms. Early-life low-protein diet induced a higher level of H3 Lys27 trimethylation (H3K27me3) by upregulation of the protein level of E(z), H3K27 specific methyltransferase. This altered H3K27me3 was associated with a short lifespan in the offspring (Xia et al., 2016). Two days of high/low-sugar diet elicited obesity in offspring, which involves H3K9/K27me3-dependent reprogramming of metabolic genes (Öst et al., 2014). However, how the nutritional stress induces epigenetic modification to be transmitted to the next generation remains unknown.

*Sir*2/sirtuin1, which encodes a NAD-dependent class III histone deacetylase is a nutrition sensor and an effective regulator of metabolism and stress responses (Imai et al., 2002). Sir2 activity depends on the cellular level of NAD+, which is increased by starvation. Activated Sir2 has many functions to maintain energy homeostasis, regulating insulin signaling, fat mobilization, and energy consumption (Banerjee et al., 2013; Banerjee et al., 2017; Palu & Thummel, 2016). Moreover, Sir2 reduction is related to dysfunction of lipid metabolism in offspring by the maternal dietary intervention (Nguyen et al., 2017). On the other hand, Sir2 overexpression in offspring attenuates intergenerational effects (Nguyen et al., 2018). These studies have suggested that Sir2 may respond to nutritional stress and lead to the generation of epigenetic modification in the parental body.

In this study, we investigated the role of *Sir2* in the intergenerational effects caused by paternal dietary conditions. We found that *Sir2* mutant males showed no intergenerational phenotypes such as an increase of TAG levels in their offspring. The RNAi-mediated knockdown of *Sir2* in the fat body of father impaired the ability to induce intergenerational phenotype. This inability was rescued by overexpression of *Sir2* in the fat body. Thus, our results indicated that that *Sir2* is essential in the fat body for fathers to induce intergenerational effects of low nutrient diet on offspring physiology.

## Results

### Paternal low nutrition diet-induced intergenerational phenotypes

To induce intergenerational effects with paternal diet, 4-to 5-day-old male flies were kept on a low nutrient diet for 2 days, and allowed to mate with virgin females which have been maintained on a control diet. After 1 day, the parents were removed, and the offspring were left to develop on a control diet. Adult male offspring were transferred to a new control diet, and 4-to 5-day-old males were weighed and used to measure the amounts of TAG levels in the whole body. The amount of TAG in the offspring was increased when fathers were maintained on a low nutrition diet for 2 days (Fig. 1A). The dietary intervention in fathers had no significant effect on body weight of the offspring (Fig. 1B). Although the increase of TAG levels was reproducibly observed, we found that the weight-normalized TAG level was more consistent compared to TAG alone (Fig. 1C). We use TAG/BW values as intergenerational phenotypes induced by paternal low nutrition diet.

**Figure 1.**
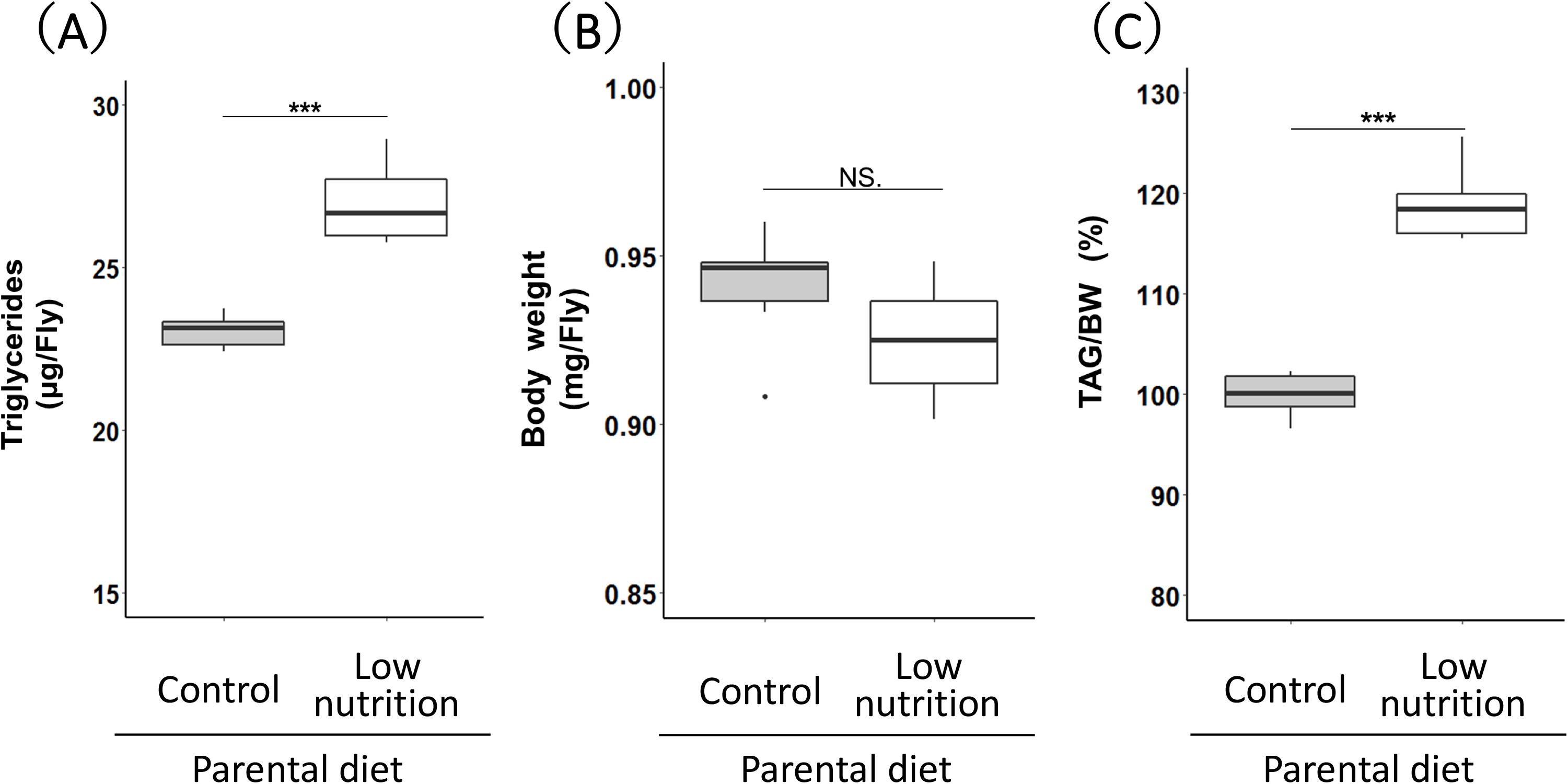
Intergenerational effects of paternal low nutrition diet on TAG levels and body weights of offspring. Triacylglyceride (TAG) levels (A) and body weight (B) and TAG/BW (C) in F_1_ males from fathers maintained on a standard diet (control) or a low nutrition diet (white) for 2 days (n = 6). Data were represented by box and whisker plot. The horizontal lines in the boxes represent the median values. Box edges and whiskers indicate the 25th/75th and 2.5th/97.5th percentiles, respectively. ***p < 0.001, NS., not significant (Student’s t-test).

Since the paternal nutrient condition could affect the physiology of offspring in various aspects, we examined whether a paternal low nutrient diet affects starvation tolerance and lifespan of the offspring flies. The paternal intervention led to increased starvation tolerance (Fig. 2A). On the other hand, the lifespan was decreased by a low diet of the fathers (Fig. 2B). These results suggested that paternal low nutrient diet has intergenerational effects on a wide variety of physiology of offspring.

**Figure 2.**
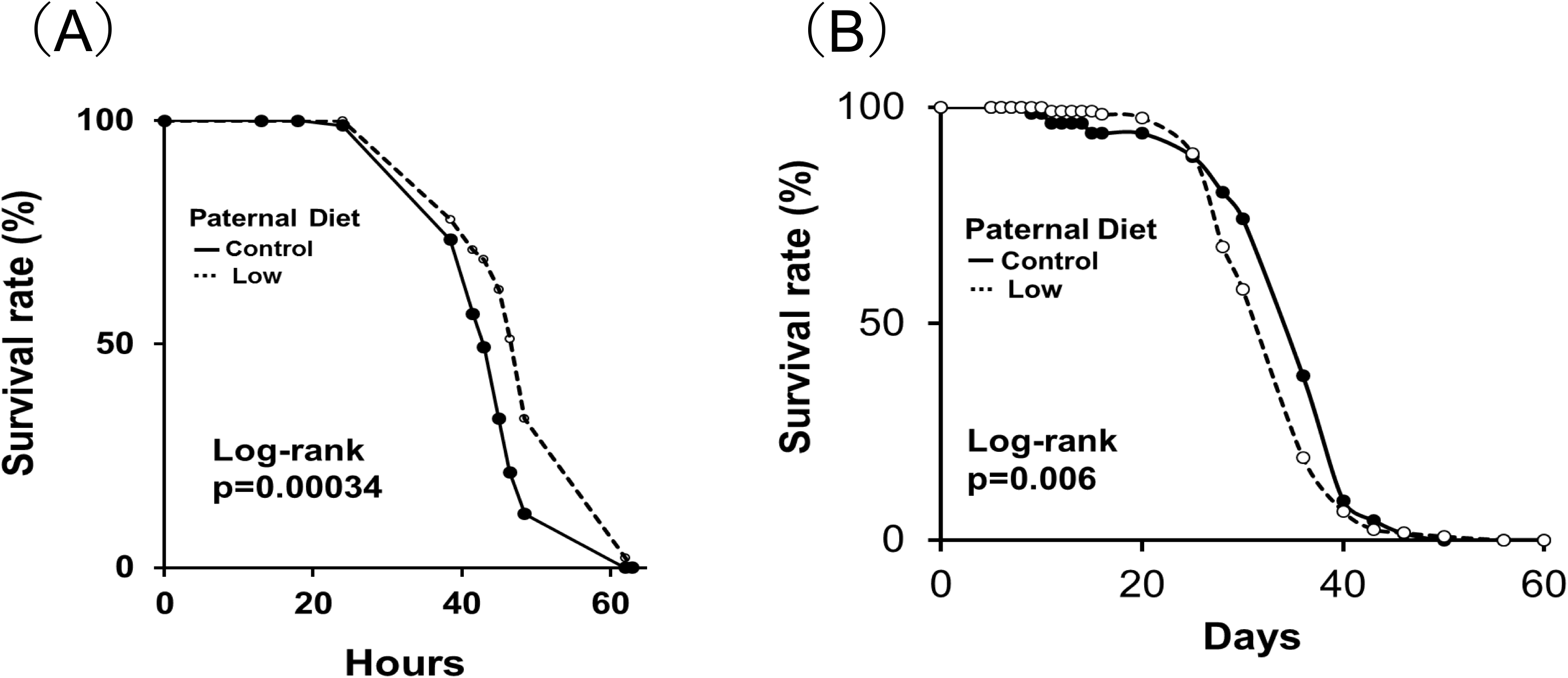
Intergenerational effects of paternal low nutrition diet on starvation tolerance and lifespan of offspring. Survival curves of F_1_ males from fathers maintained on a standard diet as a control (solid lines) or on a low nutrition diet (dotted lines) for 2 days (n = 80 for starvation tolerance assay, n = 120 for lifespan assay). Survival curves were compared between the two groups using the log-rank test. P values were indicated.

### Paternal low nutrition diet altered offspring metabolic phenotype

It has been shown that obesity and lifespan are often associated with impaired energy metabolism. Therefore, we examined whether the intergenerational effects of the paternal dietary intervention were also associated with altered energy metabolism. Using a liquid chromatography-mass spectrometry (LC-MS/MS), we measured the levels of metabolites, especially those in the primary energy-producing pathways, such as glycolysis and TCA cycle. Metabolite profiles were obtained from the offspring of fathers which were maintained on a low nutrition diet or standard diet (control) for 2 days. To compare metabolite profiles of two groups, we performed an unsupervised method of Principal Component Analysis (PCA) after the data had been preprocessed by autoscaling. The scores plot of the PCA indicated that the PC1 scores of the control and low paternal diet were negative and positive, respectively (Supplementary Fig. 1A). The PC1 score appears to be related to the diet.

Next, we investigated critical metabolic pathways related to PC1 score. We performed statistical hypothesis testing for factor loading in PC1 (Yamamoto *et al*., 2014), and 24 metabolites were statistically significant(Correlation coefficient: R ≧ 0.7, p: Holm’s method). Pathway analysis was performed for these 22 metabolites. Tricarboxylic acid (TCA) cycle, glyoxylate, and dicarboxylate metabolism and glycolysis were significantly affected in the offspring of fathers fed with low nutrition diet (Supplementary Fig. 1B). Our analysis revealed that low paternal diet altered offspring metabolic states.

### Paternal Sir2 is essential in the fat body for generating intergenerational effects

To identify a gene required for generating the intergenerational phenotype in fathers, we focused on *Sir2*, which encodes a NAD-dependent class III histone deacetylase. *Sir2* is an evolutionarily conserved nutrient sensor, which is activated by starvation or low nutrient conditions (Brachmann *et al*., 1995; Frye, 2000). Moreover, Sir2 can regulate levels of H3K9/K27me3 (Furuyama *et al.*, 2004; Vaquero *et al.*, 2007). Therefore, it is possible that Sir2 is involved in the intergenerational effects by transmitting information from fathers to offspring via germline cells. The existing evidence suggests that Sir2 may have functions to mediate low-nutrient diet-induced intergenerational effects.

To examine the role of Sir2 in fathers, we determined a weight-normalized TAG level and starvation tolerance in the offspring. The offspring of *Sir2* mutant fathers showed no TAG accumulation (Fig. 3A) and decreased starvation tolerance (Fig. 3B). Next, we performed the RNAi-mediated knockdown experiments. The knockdown of *Sir2* in the whole body showed the same phenotype as *Sir2* null mutant (Fig. 4A). Interestingly, the knockdown of *Sir2* in the fat body showed no response in the offspring (Fig. 4A). These results suggested that parental *Sir2* is required in the fat body to form intergenerational effects. To confirm that, we conducted the fat body-specific *Sir2* overexpression experiment in *Sir2* null mutant background. The TAG accumulation in the offspring was rescued by fat-body-specific *Sir2* overexpression in the *Sir2* null mutant fathers (Fig. 4B). The findings suggested that parental *Sir2* is essential in the fat body for generating intergenerational effects.

**Figure 3.**
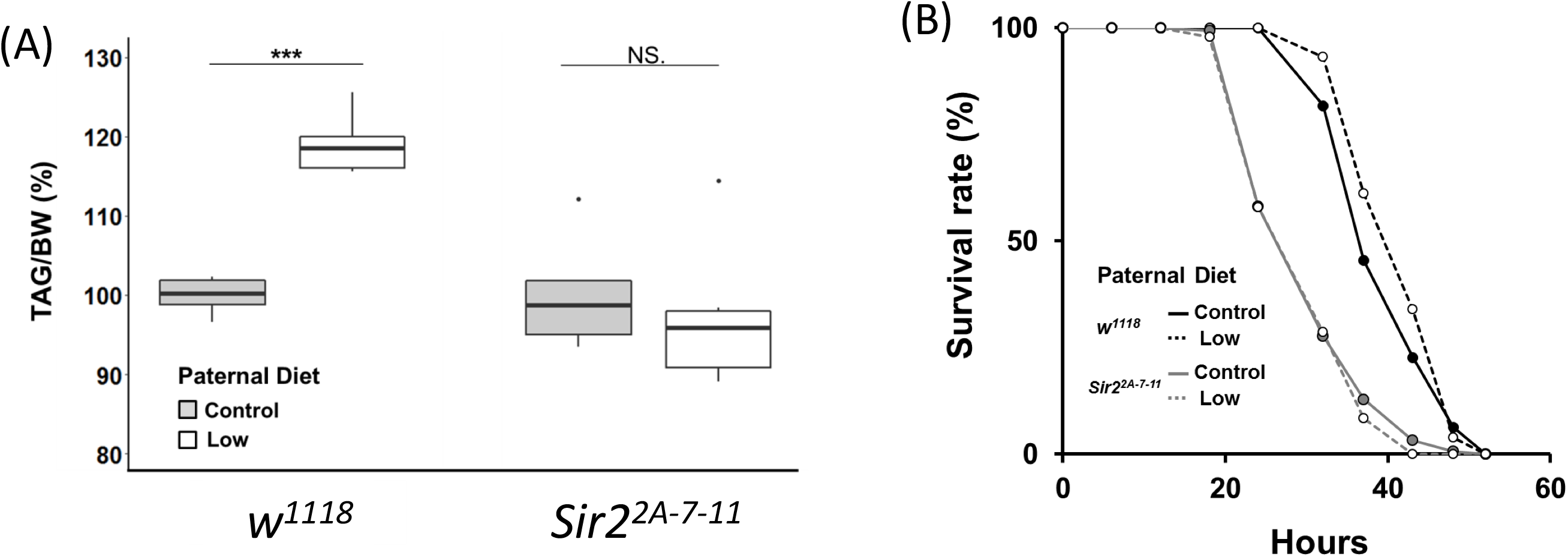
Sir2 is essential for the intergenerational effects of paternal low nutrition diet. (A) Sir2 mutants were examined for nutrition-dependent intergenerational effects using *w*^*1118*^ as a control and *Sir2*^*2A-7-11*^, a null allele. Fathers with either genotype were fed with standard diet group (black) or a low nutrition diet group (white), and TAG/BW values of offspring were determined (n = 4 for each group). (B) Starvation tolerance of F_1_ males of *w*^*1118*^ (circle) and *Sir2* mutant (diamond) fathers maintained on a control diet (solid lines) and a low diet (dotted lines) for 2 days (n = 80 for each group). P-values for the log-rank test were indicated.

**Figure 4.**
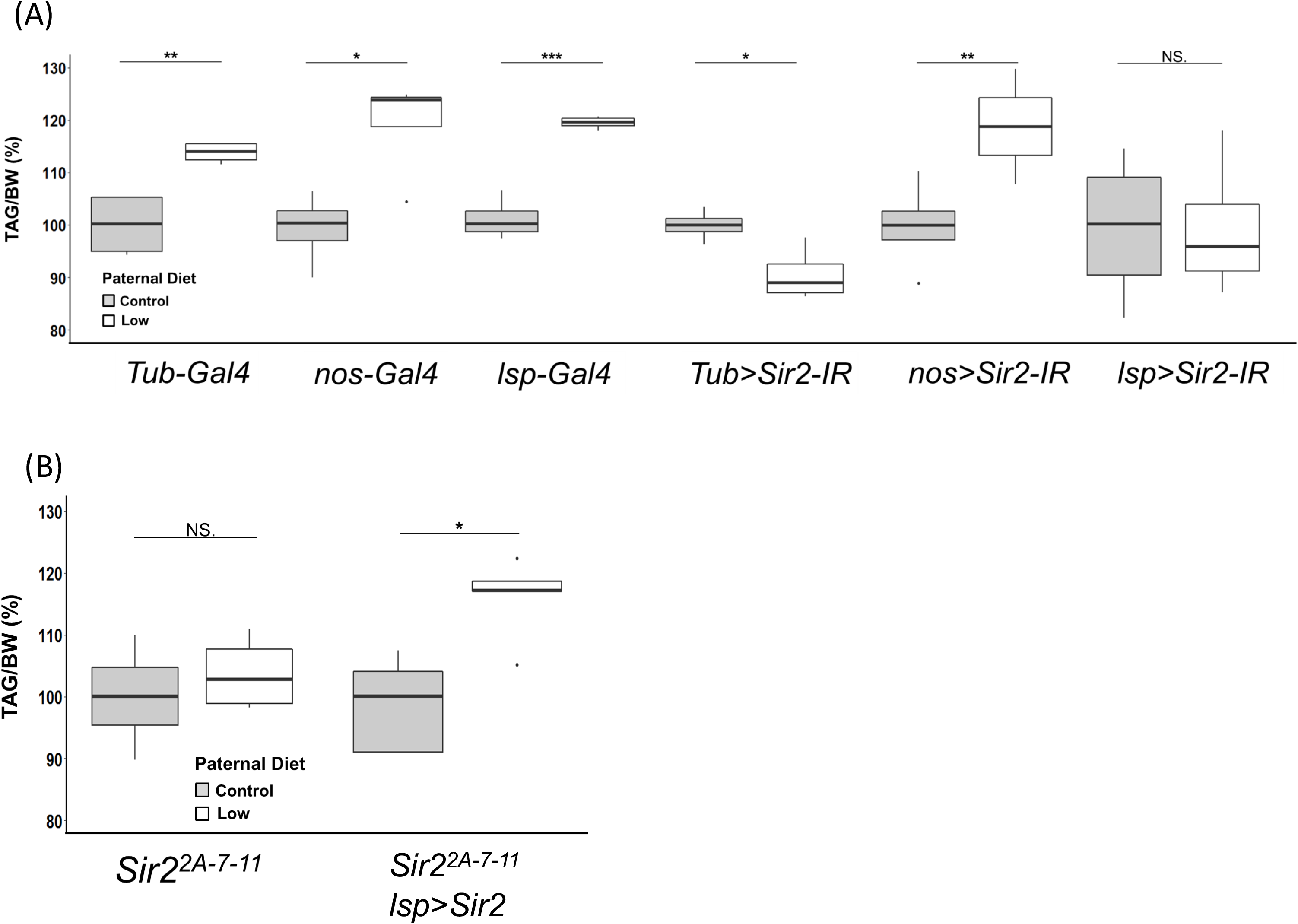
The sir2 function is critical in the fat body of fathers to induce intergenerational effects. (A) Tissue-specific knockdown of *Sir2* was performed in fathers using *UAS-Sir-IR* and various GAL4 drivers, and TAG/BW levels in the offspring were determined (white). *Tub-Gal4*, *nos-Gal4*, and *lsp-Gal4* strains were used as control genotypes of fathers. Fathers were fed with a standard diet (black) or low nutrition diet (white) for 2 days (n = 4 for each group). Intergenerational effects of low nutrition diet were observed for all control genotypes, and *nos > Sir2-IR* (Germline cells). However, TAG/BW levels did not increase in the offspring of *Tub > Sir2-IR* (whole body) and *lsp > Sir2-IR* (Fat body). (B) Overexpression of Sir2 induces intergenerational effects of low nutrition diet in Sir2 mutants. The accumulation of TAG was observed in the offspring of the *Sir2* null mutant fathers with fat-body-specific *Sir2* overexpression (n = 6 for each group). The lines in the boxes represent the medians, and box edges and whiskers indicate the 25th/75th and 2.5th/97.5th percentiles, respectively. *: p < 0.05, **: p < 0.01, ***: p < 0.001, NS., not significant (Student’s t-test).

## Discussion

In the present study, we demonstrated that low nutrition diet for fathers induced intergenerational phenotypes, which include an increased amount of TAG in the offspring. The offspring also showed reduced life span, starvation tolerance, and altered energy metabolic pathways including glycolysis, TCA cycle, glyoxylate, and dicarboxylate metabolism. Furthermore, we found that loss of paternal Sir2 showed no TAG accumulation in the offspring. The same phenotype was observed when paternal Sir2 was knocked down in the fat body, and transgenic overexpression of Sir2 in the fat body of father was sufficient to rescue mutant phenotype. These results support our hypothesis that paternal Sir2 receives nutritional stress and plays an essential role in generating information leading to transmission of intergenerational effects in the paternal body.

Previous studies have suggested that the nutrient-induced intergenerational effects are transmitted to offspring by H3K9/K27me3 in germline cells (Guida *et al*., 2019; Öst *et al.*, 2014; Xia *et al.*, 2016). Recently, it has been reported that maternal H3K9me3 is directly inherited to next generation and required for normal embryogenesis in mice and *Drosophila* (Inoue *et al.*, 2017a; Inoue *et al.*, 2017; Zenk *et al.*, 2017). However, it remains unclear how environmental signals control histone modification enzymes which produce H3K9/K27me3. It has not been clear whether environmental signals directly regulate histone modification enzymes in germline cells to induce environment-dependent intergenerational epigenetic changes. In our experiments, knockdown of *Sir2* in the germline cells (*nos* > *Sir2-IR*) did not affect the intergenerational phenotype. Although germline cells are likely to have epigenetic modifications in response to low nutrition diet, they don’t seem to respond to the signal directly. Sir2 is responsible for receiving environmental stress and may generate signals that could cause epigenetic modification in the germline cells to alter the metabolic phenotype. It is crucial to identify the downstream components of Sir2, which are involved in the transmission of environmental stress information to germline cells.

Transgenerational inheritance has been observed in various organisms. In human, many studies have reported that parental and early-life nutritional conditions have effects throughout one’s life (Langley-Evans, 2015; Tarry-adkins & Ozanne, 2011). It should be determined how intergenerational effects are generated from parental nutritional conditions in parents. Although the process could be complicated, the discovery of Si2 as a critical regulator of intergenerational effects of low nutrition diet would simplify the approach to understanding the mechanism of epigenetic inheritance.

## Experimental procedures

### Fly strains and media

Flies were maintained on a standard glucose-yeast-agar medium containing 10% glucose, 4% dry yeast, 9% cornmeal, 0.8% agar, 0.3% propionic acid and 1% para-hydroxybenzoate in temperature-controlled environmental chambers at 25°C throughout development. Unless otherwise stated, the standard medium was used as a control medium in all experiments. Low nutrition diet was prepared by diluting the standard medium (1/5), and mixed with agar at the final concentration of 0.8%. *Sir2^2A-7-11^* (Xie & Golic, 2004), *nos-Gal4* and *lsp-Gal4* were obtained from the Bloomington Stock Center. *Ptc-Gal4* and *Tub-Gal4* were obtained from the Kyoto Stock Center. The *Sir2^RNAi^* line was obtained from the National Institute of Genetics. Since *Sir2* mutant strain has *w^1118^* background, we used *w^1118^* flies as a genotype control.

### Generation of transgenic fly

To generate transgenic flies expressing *Sir2* in germline and somatic cells, we constructed pUASp-Sir2-attB using pUASp-attB vector (Kaneuchi *et al.*, 2015) and introduced them into the fly genome using a phiC31-based integration system (Bischof *et al.*, 2007).

### Triacylglyceride (TAG) measurement

TAG content was determined by an enzymatic assay with lipase (Triglyceride Reagent, Wako), which breaks down the triacylglycerol to free glycerol and three fatty acids. Ten flies (3-to 5-day-old) were weighed and homogenized with 550 µl of 1% Triton X-100 (Wako). Homogenates were incubated for 10 min at 70°C and frozen at −80°C. The frozen samples were thawed on ice and centrifuged for 10 min at 15,000 g at room temperature. The supernatants were collected and centrifuged again under the same conditions, and the resulting supernatants were used to measure the amount of TAG. A 10 µl sample was mixed with 100 µl assay reagent (free glycerol reagent, Wako) and 20 µl lipase reagent (triglyceride reagent, Wako), and incubated for 30 min at 37°C. The absorbance at 540 nm was measured using a plate reader (Enspire, PerkinElmer).

### Starvation tolerance

To determine the survival rate of flies under starvation, 3-to 5-day old male flies were kept in vials containing 2% agar, and the number of dead flies was counted every 4 h. At least 80 flies per paternal diet and genotype were used.

### Lifespan

Three-to five-day-old males flies were maintained in vials containing a standard glucose-yeast-agar medium at 25°C (30 males per vial). Flies were transferred to fresh food every 3 days, and the number of dead flies was counted at the time of transfer.

### Measurement of metabolites

Ten adult flies (3-to 5-day old) were homogenized in 200 ml of 75% acetonitrile on ice. Supernatants were collected after centrifugation for 10 min at 12,000 *g* and were dried in a miVac Sample Concentrator (Genevac, Stone Ridge, NY). The sample was resuspended in 10 mM dibutyl ammonium acetate (DBAA, pH 4.95). The metabolites were analyzed by using a liquid chromatography quadrupole time-of-flight mass spectrometry (LC-QTofMS) system, ACQUITY UPLC, and Xevo QTofMS (Waters, Milford, MA).

### Statistical analysis

Analyses of metabolites data were carried out with MetaboAnalyst 4.0 (http://www.metaboanalyst.ca), a web-based platform for metabolomics data analysis (Chong et al., 2018). The data were initially analyzed using the unsupervised method of principal component analysis (PCA), and then further analyzed using statistical hypothesis testing for factor loading in PC1 (statistically significant: Correlation coefficient: R ≧ 0.7, p: Holm’s method). Pathway analysis for factor loading was performed independently for metabolites with a significant difference between the two groups.

Statistical significances are evaluated using Student’s t-test for comparison of the levels of TAG and metabolites. The log-rank test was used for comparison of survival curves. A Box and Whisker Plot was used to visualize the data of TAG levels, body weights, and TAG levels normalized by body weights. The middle lines in the boxes represent the medians. Box edges and whiskers indicate the 25th/75th and 2.5th/97.5th percentiles, respectively.

## Acknowledgments

We are grateful to the members of the Cellular Genetics Lab for helpful discussions and critical reading of the manuscript. We also thank the Kyoto Stock Center, the Bloomington Stock Center, and the National Institute of Genetics for providing the stocks.

---

**Supplementary Figure 1.**
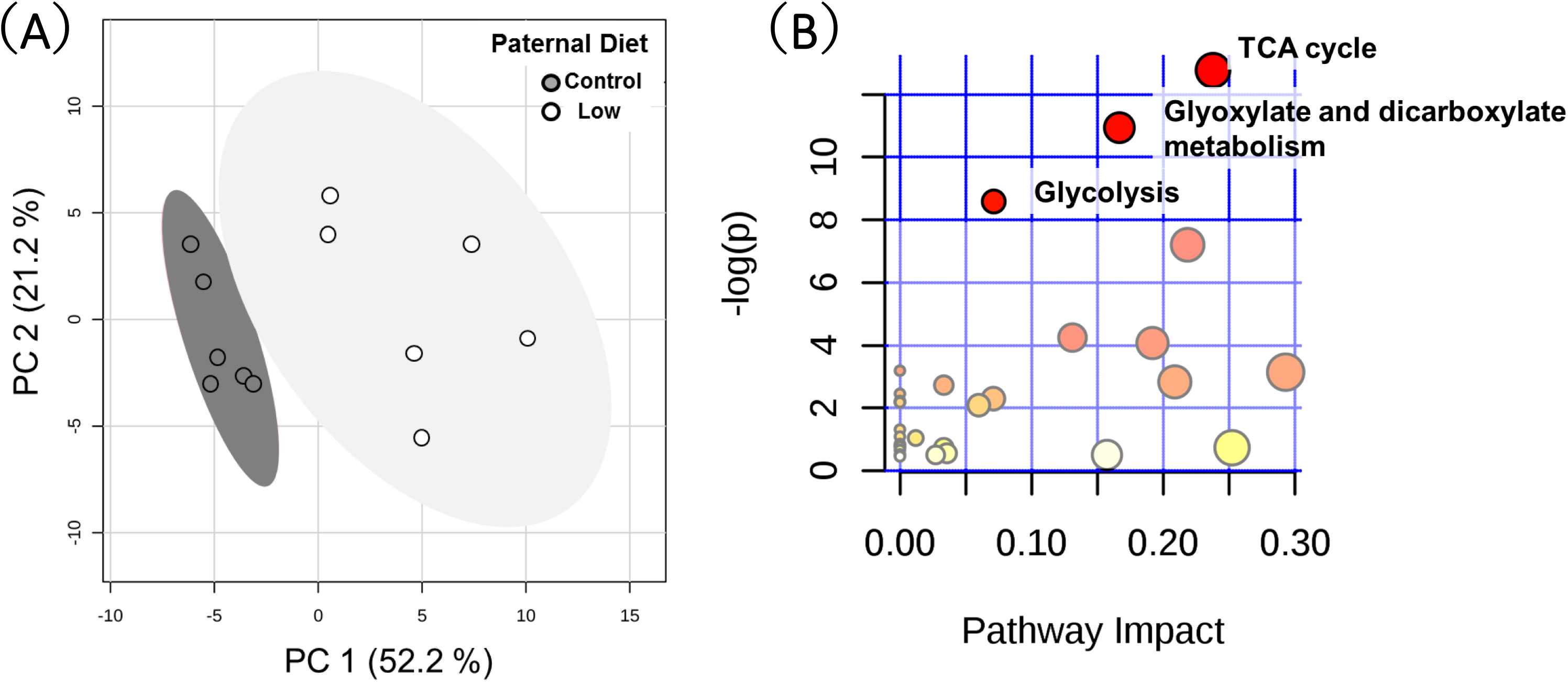
Low paternal diet altered offspring metabolic phenotype. (A) Principal component analysis (PCA) of metabolite data from F_1_ males of fathers maintained on a control diet (black circle) and those fed with a low diet for 2 days (white circle). The first principal component accounts for 52.2% and the second principal component accounts for 21.2% of the variance. (B) Metabolic pathways affected by low nutrition diet. *P* values were calculated using MetaboAnalyst 4.0 software (Chong et al., 2018).

